# Nanofluid-Enhanced Laser Lithotripsy Using Conducting Polymer Nanoparticles

**DOI:** 10.1101/2024.06.01.596977

**Authors:** Qingsong Fan, Junqin Chen, Arpit Mishra, Aaron Stewart, Faisal Anees, Ting-Hsuan Chen, Judith Dominguez, Christine Payne, Michael E. Lipkin, Pei Zhong, Po-Chun Hsu

## Abstract

Urinary stone disease, characterized by the formation of hard mineral deposits in the urinary tract, has seen a rising prevalence in the U.S. in recent years. This condition often leads to severe pain and typically requires medical intervention. Laser lithotripsy, a minimally invasive treatment, uses laser energy to fragment urinary stones into smaller pieces, facilitating easier removal or natural passage. Among available laser technologies, the holmium:yttrium-aluminum-garnet (Ho:YAG) laser has established itself as the gold standard over the past three decades. Efforts to improve Ho:YAG laser ablation efficiency have largely focused on adjusting laser parameters such as pulse energy and frequency. In this study, we proposed a nanoplasmonic engineering strategy by incorporating nanoparticles (NPs) with strong near-infrared (NIR) absorption into the fluid surrounding the stone, enhancing the light-matter interaction. Using a 0.03 wt.% PEDOT:PSS nanofluid, stone ablation efficiency improved by 38–727% in spot treatment and 26–75% in scanning treatment with a clinical laser lithotripter. The highly absorbing nanofluid accelerates vapor tunnel formation, boosts laser energy transmission to the stone, and permeates stone pores to enhance damage, without increasing thermal tissue injury risk. Cytotoxicity tests also confirmed minimal toxicity at appropriate concentrations. This nanofluid-based approach offers a promising advancement for more efficient and safer laser lithotripsy.

## Introduction

Urinary stone disease (USD) is a benign yet severely painful genitourinary condition affecting nearly 1 in 10 Americans.^[1]^ In 2000, the annual health expenditure for USD in the U.S. exceeded $2 billion, which continues to rise rapidly today.^[2–5]^ For USD patients, minerals such as calcium oxalate, calcium phosphate, and uric acid, gradually crystallize and form large stones that cannot be naturally expelled from the urinary system. When these stones grow to a size capable of obstructing urine flow, they cause intense pain along with additional symptoms like frequent urination, difficulty urinating, and blood in the urine.^[6]^ Various techniques have been developed to remove urinary stones, including extracorporeal shock wave lithotripsy, laser lithotripsy (LL) via ureteroscopy, and percutaneous nephrolithotomy.^[7]^

LL is the most rapidly growing intervention method for USD treatment. Take extracorporeal shock wave lithotripsy for example, despite its advantages of non-invasiveness and anesthesia-free treatment, it exhibits lower efficacy in managing large (> 1 cm) or hard stones, resulting in reduced clearance and stone-free rates.^[8–10]^ In contrast, LL has made significant advancements, particularly those utilizing the holmium: yttrium-aluminum-garnet (Ho:YAG) laser,^[11,12]^ which has the suitable pulse duration, repetition frequency, and peak power to offer numerous advantages, such as efficacy in fragmenting all types of stones,^[13–15]^ minimal retropulsion,^[16]^ and compatibility with low-cost flexible optical fibers. Also importantly, large series of clinical studies have shown that Ho:YAG LL is safe in children,^[17–19]^ in all stages of pregnancy^[20]^ and in patients with bleeding disorders.^[21,22]^ All in all, Ho:YAG LL has been the gold standard in USD management.^[23,24]^

For both research and clinical practice of Ho:YAG LL over the past two decades, one primary goal is to maximize stone ablation efficiency with minimal thermal injury. From the optical science perspective, this effort could be facilitated by controlling the light-matter interaction, particularly through modulating the absorption coefficient. Nevertheless, the majority of prior research efforts have focused on manipulating the laser output profile, including pulse energy, pulse duration, and frequency.^[25–31]^ These prior approaches did not change the fundamental physical properties of water and stone at the wavelength of Ho:YAG (λ = 2120 nm), so the accessible parameter space and enhancement are limited. There were a few studies that painted the kidney stone with either laser-absorbing pigments or nanoparticles (NPs) for augmenting stone damage in an *in vitro* setting, However, the clinical applicability of this approach remains to be further explored, particularly in terms of how such a coating might be maintained or reapplied following surface ablation.^[32,33]^ In addition, it has been reported that the collapse of the vapor bubble produced at the fiber tip can play a critical role in stone dusting during LL, indicating the importance of attending to not only the stone but also the surrounding fluid environment.^[34,35]^

Here, we propose to use nanoparticle dispersion, i.e., nanofluid, to enhance the laser energy absorption and, thus, the ablation efficiency of Ho:YAG LL. In this study, we introduce poly(3,4-ethylenedioxythiophene) polystyrene sulfonate (PEDOT:PSS) NPs, a well-established polymeric material known for its absorption peak in the near-infrared spectrum, into the fluid. At a concentration of 0.03 wt.%, the PEDOT:PSS NPs can increase the absorption coefficient of the fluid at 2120 nm by 25% without compromising the visibility of the field of view in the ureteroscope. Based on the process of the stone ablation during LL (**Scheme 1b**), we hypothesized that vapor bubble can be generated and bridge the laser fiber and stone surface at the earlier stage of each laser pulse, facilitating the delivery of laser energy to the stone. In addition, the PEDOT:PSS NPs trapped within the stone or attached to the surface could increase the absorption of the transmitted laser energy, which leads to higher thermal ablation and micro-explosion. As a proof-of-concept, we explored the efficiency of PEDOT:PSS nanofluid for LL on BegoStone via spot and scanning treatment. LL conducted in a 0.03 wt.% PEDOT:PSS nanofluid demonstrated an improvement of 38 – 727% in spot treatment and 26 – 75% in scanning treatment in stone ablation compared to procedures conducted in water at the fiber tip-to-stone standoff distance (SD) of 0 – 1 mm. In addition, the minimal impact of PEDOT:PSS nanofluid at this concentration on the thermal injury and viability of mIMCD-3 cells evaluated via the hydrogel kidney model and cytotoxicity test demonstrated its potential utilization in clinical Ho:YAG LL without compromising safety.

## Results and Discussion

### Optical properties of PEDOT:PSS nanofluid

At the Ho:YAG wavelength of 2120 nm, PEDOT:PSS stands out as a promising conductive polymer-based nanofluid because of its free-carrier absorption enabled by the PEDOT delocalized hole carriers and its water dispersibility achieved by the sulphonic groups in the PSS moiety. PEDOT:PSS also finds widespread applications in flexible electronics,^[36–40]^ solar cells,^[41,42]^ IR sensors,^[43]^ and more. Its appeal lies in its tunable electrical conductivity, relative transparency to visible light, excellent thermal stability, and biocompatibility.^[44]^ Moreover, PEDOT:PSS NPs are conveniently synthesized through colloidal routes,^[45]^ facilitating large-scale production at a relatively low cost. As depicted in **Figure 1a**, the addition of PEDOT:PSS NPs to water leads to a gradual increase in fluid absorbance within the NIR range. Of particular interest is the absorbance of fluids at 2120 nm, the operational wavelength of the Ho:YAG laser. By plotting the absorbance at 2120 nm against the concentrations of PEDOT:PSS nanofluid, a linear correlation emerges between these parameters, showing the typical Beer’s law behavior (**Figure 1b**). Since the penetration depth is the inverse of absorption coefficient, it becomes shorter as the concentration of the PEDOT:PSS nanofluid increases. For example, the penetration depth of light at 2120 nm diminishes from 379.6 μm for water to 342.2 μm for 0.01 wt.% and further to 285.3 μm for the 0.03 wt.% PEDOT:PSS nanofluid. These decreases of penetration depth roughly correspond to a 10% and 25% increase of absorbed power density.

**Scheme 1.**
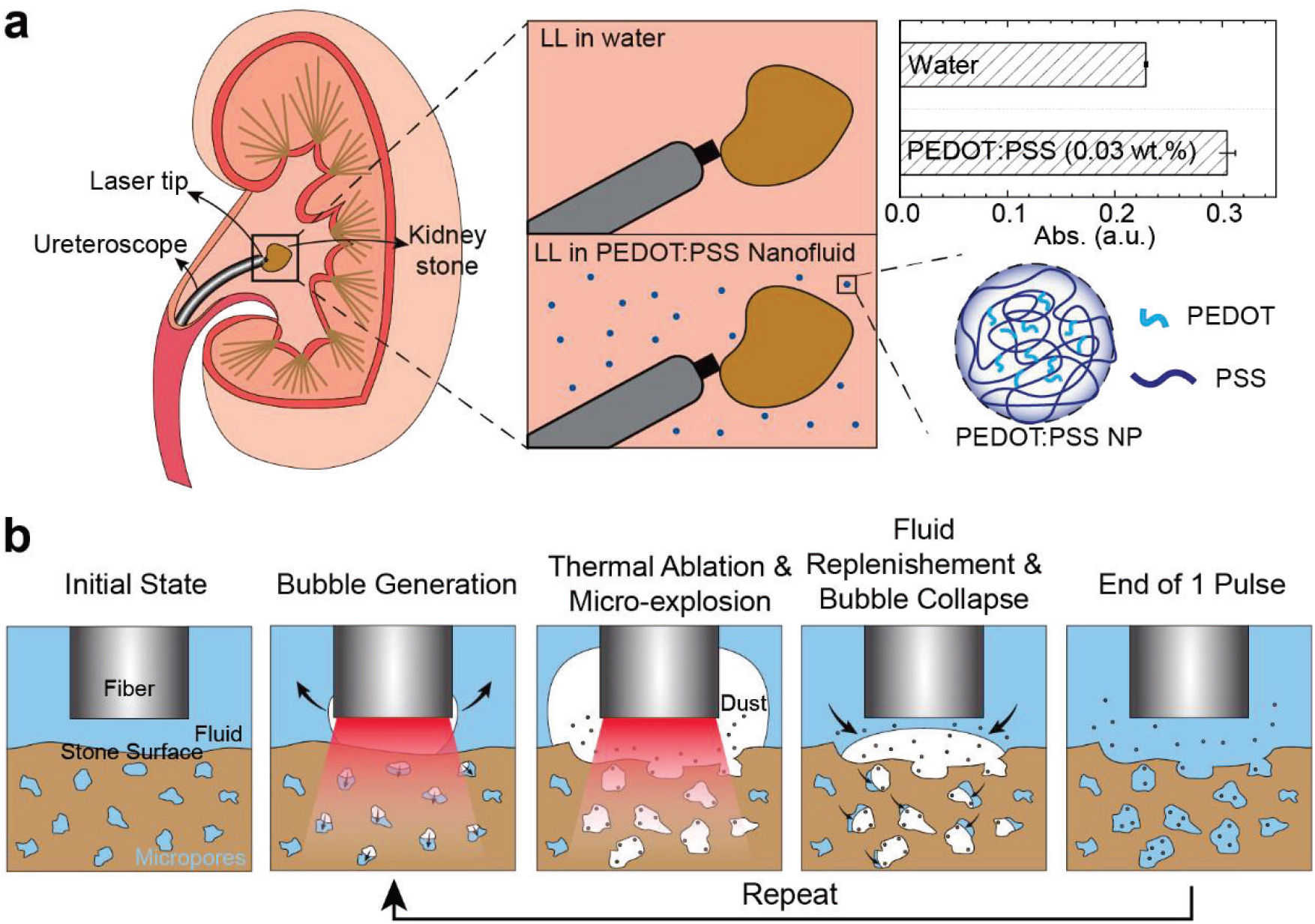
Schematic illustration. (a) LL in kidney stone removal. Traditional LL is conducted in saline (water) with a relatively low absorption coefficient at 2120 nm. Introducing PEDOT:PSS nanofluid can enhance the stone damage efficiency by increasing the absorption coefficient. (b) Mechanisms of stone ablation during LL. Thermal ablation due to the absorption of laser energy by the stone surface, micro-explosion of the trapped fluid inside the stone, and violent bubble collapse all contribute to the stone damage.

**Figure 1.**
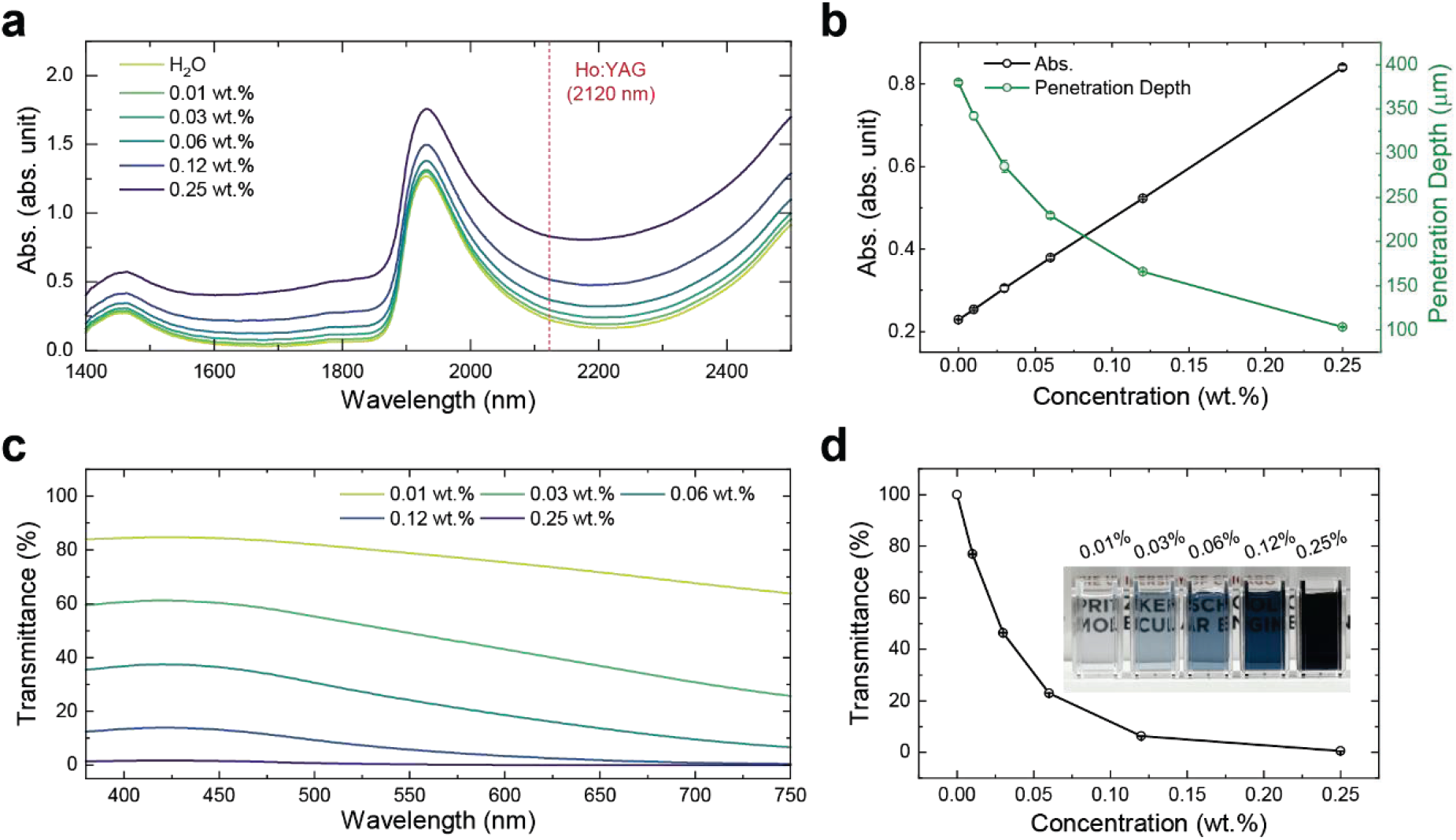
Optical properties of PEDOT:PSS nanofluid. (a) Absorbance spectra of water and PEDOT:PSS nanofluids of different concentrations within the NIR range (path length of measurements: 0.2 mm). (b) The absorbance and derived penetration depth values at the wavelength of 2120 nm of PEDOT:PSS nanofluids. (c) Transmittance of PEDOT:PSS nanofluids in the visible spectrum (path length of measurements: 5 mm). (d) The calculated average transmittance over a wavelength range of 380-750 nm (inset: a digital photo of PEDOT:PSS nanofluids at different concentrations in cuvettes with a 5 mm path length).

Although adding more PEDOT:PSS NPs in water can continuously enhance the NIR absorption, it has an upper limit posed by the reduced visible clarity, a crucial factor for the urologists to locate the stone and laser spot. The π-π* transition^[46]^ of PEDOT:PSS results in the absorption of visible light that makes the solution appear dark blue, especially at high concentrations (**Figure 1d**, inset). This strong visible absorption contradicts with the stringent requirement of a clear field of view for the surgeon during the LL. To determine the optimal concentration of PEDOT:PSS nanofluid for LL by considering both NIR absorbance enhancement and visibility, we measured the transmittance of the above solutions within the visible regime (**Figure 1c**). The path length was chosen to be 5 mm to mimic the typical searching and working distance between the ureteroscope and the kidney stone in clinical practice.^[47]^ The average visible transmittance of the nanofluids was found to decrease with increasing PEDOT:PSS concentration, as expected. Upon inspecting the field of view under the ureteroscope, we observed that the kidney stone remains clearly visible with the concentration lower than 0.03 wt.% (**Figure S1**).

### Hypothesis of nanofluid-enhanced stone ablation

In clinical practice, LL is conducted on stones surrounded by saline, whose primary component is water with a relatively low absorption coefficient for 2120 nm Ho:YAG laser. The well-recognized explanation to the stone damage during LL is demonstrated in **Scheme 1b**: When the pulsed laser is activated, the fluid around the fiber tip absorbs the photon energy, which turns into heat. Once the heat exceeds vaporization enthalpy (latent heat), vapor bubbles will form and expand. The formation of vapor bubbles is critical for the destruction of urinary stones for two reasons: (1) once the vapor bubble bridges the gap between the laser fiber tip and stone surface, the photon energy can be transmitted to the stone with minimal loss since water vapor has a much lower absorption coefficient compared to its liquid state, which is known as the Moses effect,^[27]^ and (2) the violent bubble collapse near the stone surface can induce mechanical damage (cavitation) to the stone^[48]^ other than conventional photothermal ablation theory for laser-material interactions.

We hypothesized that by increasing the absorption coefficient of the surrounding fluid, more photon energy will be absorbed to generate the vapor bubble at the beginning of the laser irradiation, leading to a faster vapor tunnel establishment and therefore more photon energy delivered to the stone surface. To verify this, we investigated the change of bubble behavior with the addition of PEDOT:PSS NPs, especially the size and expansion rate of the bubbles, using a high-speed camera (Phantom v7.3, Vision Research, Wayne, NJ) operating at 100,000 frames per second. **Figures 2a** and **2b** depict the initial frames of bubble generation and expansion in different PEDOT:PSS concentrations at standoff distances (SDs) of 0.5 mm and 1 mm, respectively. To reproduce the laser-fluid-stone interaction during LL, a glass slide was placed in front of the fiber at a certain distance. After measuring the distance between the fiber tip and the bubble apex in each frame (**Figure 2c** and **Supplementary Note S2**), we found that the bubble in 0.03 wt.% PEDOT:PSS nanofluid indeed expanded the fastest at both SDs. For example, compared to that in water, it took ∼10 μs and ∼30 μs less for the bubble apex to reach the distance of 0.5 mm and 1 mm in 0.03 wt.% PEDOT:PSS nanofluid, respectively. We also measured the surface tension of PEDOT:PSS nanofluids up to the concentration of 0.25 wt.%, which remained constant (**Figure S3**). This finding demonstrates that the faster expansion of vapor bubbles in PEDOT:PSS nanofluid is solely due to its enhanced NIR absorbance, indicating a distinctly different mechanism from previous work.^[49,50]^

**Figure 2.**
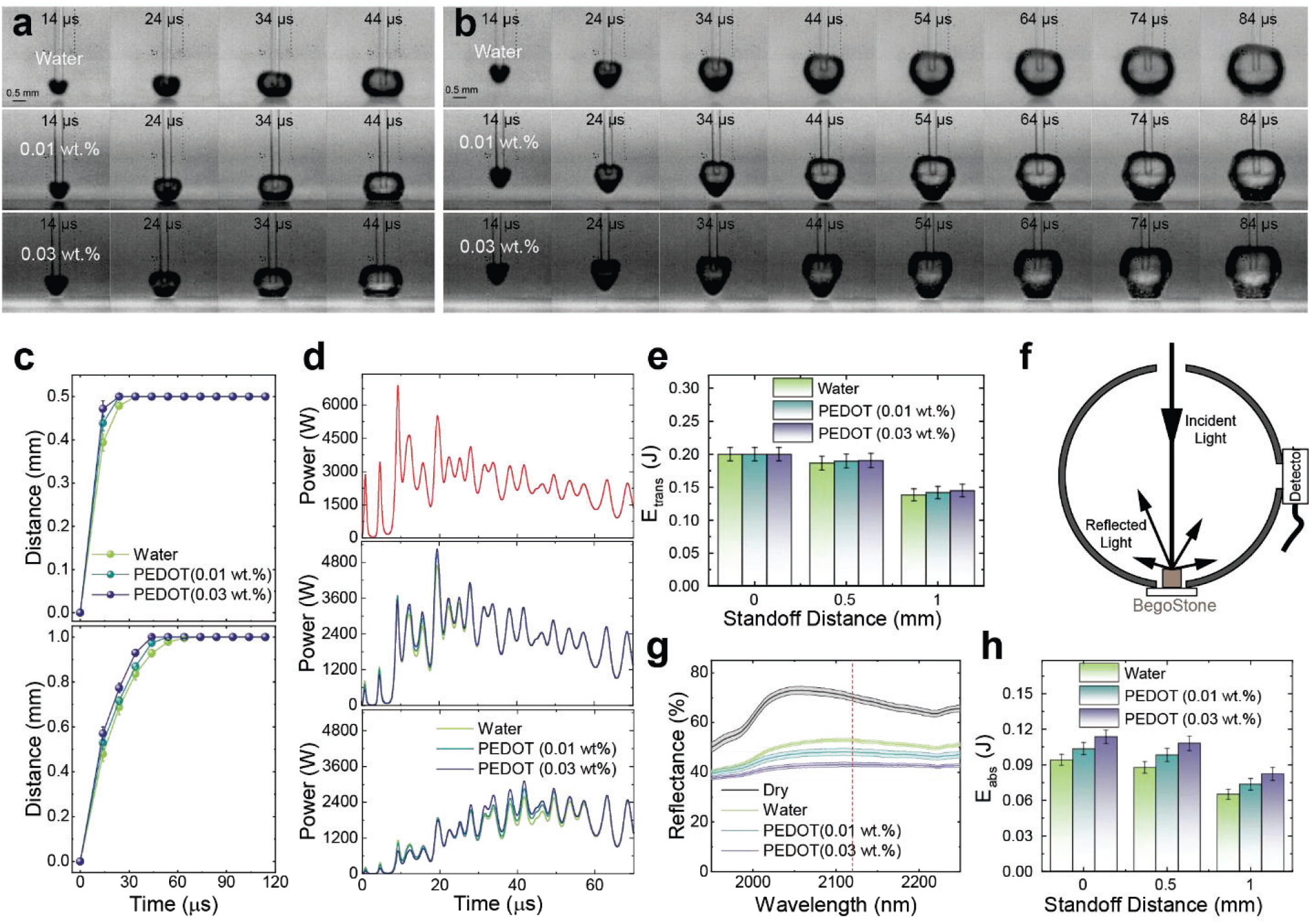
Bubble dynamics and calculated photon energy. (a, b) Snapshots of bubbles generation and expansion during one pulse of the LL in different fluids (from top to bottom: water, 0.01 wt.%, and 0.03 wt.% PEDOT:PSS nanofluid) with a glass slide placed at a distance of 0.5 mm (a) and 1 mm (b) in front of the laser fiber. (c) Temporal change of the distance between fiber and bubble apex in various fluids (top: SD = 0.5 mm, bottom: SD = 1 mm. Error bars are standard deviations of 45 measurements). (d) The output power of the 60^th^ pulse of the Ho:YAG laser (top) and calculated transmitted powers in different fluids (middle: SD = 0.5 mm, bottom: SD = 1 mm). (e) Calculated average transmitted energy at different SDs (Error bars are standard deviations of 45 measurements). (f) Scheme of NIR reflectance measurement of the soaked stones using an integrating sphere. (g) Measured NIR reflectance of stones. (h) Calculated energy absorbed by the stones soaked with various fluids.

Furthermore, we calculated the transient transmittance of 2120 nm light by considering the total optical density along the light path (**Figure S4**). By multiplying the transient transmittance with the laser output power (**Figure 2d**, top), we obtained the transient transmitted powers at these SDs (**Figure 2d**, middle and bottom). Higher laser powers could be transmitted at the early stage of each pulse due to the faster establishment of the vapor tunnel in 0.03 wt.% PEDOT:PSS nanofluid. The transmitted energy was then calculated by integrating the transmitted power over time. In **Figure 2e**, we assumed that the laser energy of each pulse could be transmitted with no loss (0.2 J) at the SD of 0 mm. At the SD of 1 mm, there was a 4.8% increase in transmitted energy for 0.03 wt.% PEDOT:PSS compared to water. Similarly, this increase was 2.1% at the SD of 0.5 mm.

In addition to the faster vapor tunnel establishment, we speculated that nanofluid may also influence the stone behavior, especially its ability to absorb transmitted photon energy. It has been widely recognized that BegoStone and human kidney stones contain numerous submillimeter pores,^[51]^ allowing the surrounding fluid to percolate into the stone and occupy the small pores. Additionally, NPs such as PEDOT:PSS may attach to the stone surface due to chemical or charge-induced adsorption. As a result, both PEDOT:PSS NPs trapped in the pores and those attached to the stone’s surface contribute to increased absorption of laser energy. We measured the NIR reflectance of BegoStones with various trapped fluids using a spectrometer equipped with an integrating sphere (**Figure 2f**), and the results are summarized in **Figure 2g**. As anticipated, stone reflectance at 2120 nm decreased from 52.9% to 43.2% when water was replaced with 0.03 wt.% PEDOT nanofluid. We further derived the absorbed energy by considering both transmitted energy and BegoStone absorption when soaked with fluids. **Figure 2h** illustrates a 20.6%, 23.2% and 26.2% increase in absorbed energy in 0.03 wt.% PEDOT:PSS nanofluid compared to water, at SDs of 0, 0.5, and 1 mm, respectively.

To further demonstrate that increasing the absorption coefficient of the fluid can help to enhance photothermal ablation, we performed the LL on the fluid-soaked stones in air, where thermal ablation is the dominant mechanism (**Figure 3a**). In clinical practice, the urologist controls laser pulse energy and pulse frequency to achieve different stone damage modes, such as “dusting”, “fragmenting”, and “pop-dusting”.^[28,52–54]^ Here, we chose to operate in the dusting mode with a low pulse energy of 0.2 J and a high pulse frequency of 20 Hz, which pulverizes the stone into very fine particles and is the most common and essential mode in clinical LL for all stone types. 50 μL of the fluid was dropped on the top surface of the BegoStone, and it was quickly absorbed within seconds without a visible liquid film on top. The laser fiber tip was placed in direct contact with the stone surface (i.e., SD = 0 mm) to maximize the photon energy delivered to the stone. **Figures 3b** and **3c** summarized the appearances and dimensional measurements of the craters produced on the BegoStones wetted by various fluids after 60 pulses treatment. There is a dramatic increase in the size of the crater when PEDOT:PSS NPs were added to the fluid. Quantitatively, compared to water, the use of 0.01 wt.% and 0.03 wt.% PEDOT:PSS nanofluid results in an enhancement of 511% and 708% in the crater’s volume, showcasing the significant advantage of enhancing the absorption coefficient of the fluid via the addition of PEDOT:PSS NPs. It is worth noting that the enhancement in ablation efficiency demonstrated in this experiment was presumably ascribed to the improved photothermal ablation, with no contribution from the cavitation (no bubble collapse).

**Figure 3.**
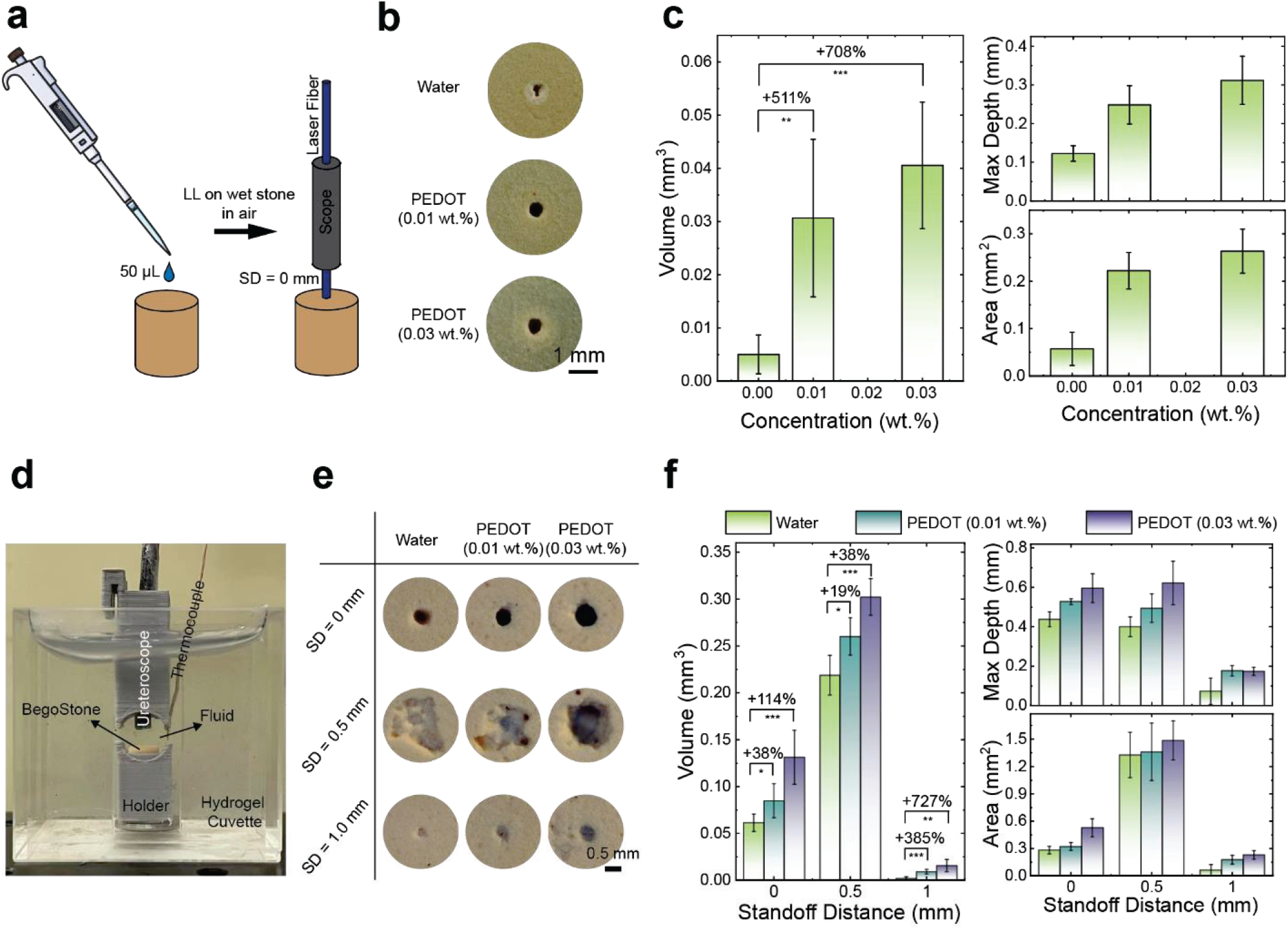
Stone damage assessment. (a) Schematic illustration of the experiment of LL in air on wet stones. (b) Photos of craters produced on BegoStone samples soaked with different fluids. (c) Dimensional measurements of craters produced on the wet stones by optical coherence tomography (OCT). (d) Experimental setup for spot treatment in fluids. (e) Photos of damage craters on BegoStone samples produced in different fluids at various standoff distances (SDs). (c) Dimensional measurements of craters on stones via spot treatment by optical coherence tomography (OCT). The values and error bars are the average results and standard deviations of 5 independent experiments under each condition. Significance of the volume measurements in (c) and (f) was calculated using Student’s test. *p<0.1, **p<0.01, ***p<0.001, ****p<0.0001, ns – not significant comparison are not presented.

### Stone damage assessment

For proof of concept, we assessed the stone damage efficiency of LL in a cuvette made of hydrogel with mechanical properties similar to the soft tissue (**Figure 3d**), containing an artificial BegoStone phantom (6×6 mm cylinders) immersed in nanofluids with three different concentrations. A ureteroscope-integrated laser fiber with an offset distance (OSD) of 3 mm was placed atop the stone at various SDs (0, 0.5, and 1.0 mm). Ideally, maximizing photothermal ablation entails direct contact between the laser fiber tip and the stone surface (i.e., SD = 0 mm). However, maintaining such contact precisely throughout the treatment is impracticable for urologists in clinical practice due to retropulsion. Therefore, we evaluated the stone damage produced in PEDOT:PSS nanofluid at three different SDs. **Figure 3e** depicts the craters produced by delivering 60 laser pulses at SD = 0, 0.5, and 1 mm in different PEDOT:PSS nanofluids, with the corresponding dimensional measurements summarized in **Figure 3f**. Craters with a circular shape were produced at SD = 0 and 1 mm, which means the photothermal ablation dominates under these conditions. However, the craters produced at SD = 0.5 mm had not only the more irregular shapes, but also the greatest volumes for all three fluids tested, showcased the additional contribution from the maximized cavitation damage, which is consistent with our previous findings.^[34,35]^ As the concentrations of PEDOT increased, the size of damage craters expanded across all SDs. Faint blue tints were consistently observed inside the craters when PEDOT:PSS nanofluids were employed. We attributed such coloration to the adherence of PEDOT:PSS NPs to the stone surface due to the local high temperatures induced by complex laser-stone interactions.

At each SD, we observed a significant increase in crater volume with the addition of PEDOT:PSS. For instance, at SD = 0 mm, the stone treatment in 0.01 wt.% and 0.03 wt.% PEDOT:PSS nanofluids resulted in crater volume increases of 38% and 114%, respectively, when compared to those produced in water under the same conditions. This enhancement in ablation efficiency was ascribed to the increase of both maximum crater depth and profile area (**Figure 3f**). Similarly, as SD increased to 1 mm, although the ablation efficiency dropped due to the attenuation of laser energy by the fluid, 0.03 wt.% PEDOT:PSS nanofluid exhibited a 727% enhancement in crater volume compared to water, unequivocally showing the unique advantages of using PEDOT:PSS nanofluids in extending the effective fiber-to-stone working distance during LL. It is worth noting that the craters produced in the fluids were much larger than those produced in air at SD = 0 mm (**Figure 2c**). We speculated that this is due to the replenishment of fluid into the pores during the off-duty cycle, making the continuous micro-explosion contributing to the stone damage throughout the treatment. Whereas for the LL in air, the trapped fluid quickly evaporated after a few pulses due to the local high temperature.

Instead of holding the laser fiber at a fixed location (spot treatment), the most common technique for dusting during the clinical practice is “painting”, where the laser fiber is moved across the stone’s surface.^[53]^ To approach this realistic clinical condition, we designed the following scanning treatment (**Figure 4a**),^[55]^ in which the laser fiber was moved at a constant speed (0.3 mm/s) across the BegoStone slab (23 x 23 x 10 mm^3^, L x W x H) during LL, with the same laser settings (0.2 J and 20 Hz) but a longer treatment duration (5 min, equivalent to 6000 pulses). Ablation efficiency was evaluated by measuring the difference in dry stone mass before and after treatment at three different SDs (0.1, 0.5, and 1 mm). The results in **Figure 4b** showcased improved ablation efficiency of PEDOT:PSS nanofluid compared to water at all three SDs: the enhancement increases from 26% to 75% as the SD increases from 0.1 mm to 1 mm. In addition, the best ablation performance, again, was obtained at SD = 0.5 mm for both water and PEDOT:PSS nanofluid, which is due to the additional contribution from cavitation that we discussed above. All these observations are consistent with the results obtained from the spot treatment (**Figure 3f**).

**Figure 4.**
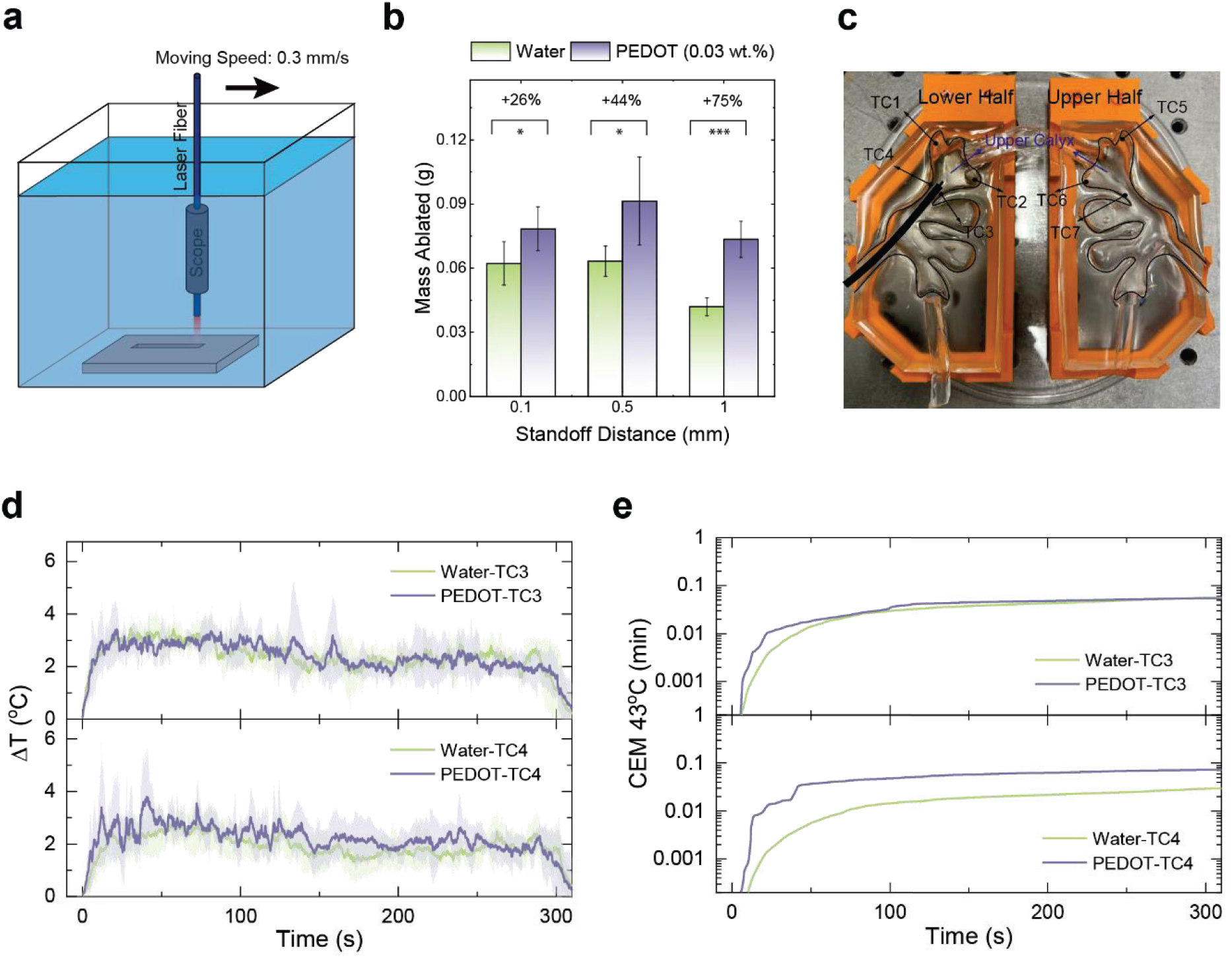
(a) Scheme illustration of the experiment setup in the scanning treatment. (b) Mass ablated from the BegoStones in scanning treatment. Significance was calculated using Student’s test. *p<0.1, ***p<0.001. (c) Geometry of the hydrogel kidney model and the locations of seven thermocouples. (d) The temperature changes of thermocouples #3 and #4 during the LL. The solid line and shaded areas are the mean and standard deviation of 5 independent measurements. (e) The corresponding CEM 43°C calculated based on the average temperature change in (d).

The temperature increase is another concern in clinical practice due to the potential thermal injury to kidney tissue.^[56–58]^ To evaluate the temperature change of the fluid in the confined cavity during the LL, we prepared a kidney model using hydrogel (**Figure 4c**). The laser fiber was inserted into the upper calyx, where seven thermocouples were located to monitor the temperature of the fluid inside, with four at the bottom half and the other three at the upper half.^[59,60]^ During LL, water or 0.03 wt.% PEDOT:PSS nanofluid at room temperature (22 ^o^C) with a flow rate of 20 mL/min was continuously flushed into the upper calyx to remove the laser-generated heat. The recorded temperature changes of the fluid during the 5-minute LL treatment at 0.2 J/20 Hz, i.e., the same laser settings used in the scanning treatment, are shown in **Figures 4d** and **S6**. Although the temperature readings from the thermocouples differed, with thermocouples #3 and #4 exhibiting the highest temperature rise, they all quickly reached a plateau after the laser was activated. Cumulative equivalent minutes at 43°C (CEM 43°C) were calculated based on the recorded temperature change to assess the potential thermal injury risk of the renal tissue (**Figures 4e and S6**). It is worth noting that the highest value of calculated CEM 43°C was only 0.073 min (Thermocouple #4 for 0.03 wt.% PEDOT:PSS nanofluid), which is far below the thermal dose threshold of kidney tissue (120 min).^[61]^ Under such a condition, the LL can be performed safely without concern for thermal injury to the renal tissue. The side-by-side comparison of stone damage efficiency and temperature rise produced in water and PEDOT:PSS nanofluid clearly demonstrated that increasing the absorption coefficient of the fluid by 25% via the addition of PEDOT:PSS NPs leads to a significant improvement of ablation efficiency by at least 26%, without compromising the safety of the procedure. Furthermore, because the stone size is finite, the significant ablation efficiency improvement using PEDOT:PSS nanofluid can potentially shorten treatment duration, reducing the risk of accumulated thermal injury and other surgical complexities.

### Cytotoxicity

PEDOT:PSS in various forms, including microwires,^[62,63]^ porous microparticles,^[64]^ films,^[65,66]^ and cell scaffolds,^[67]^ have been reported with good biocompatibility. In addition, a PEDOT-based coating (Amplicoat, Heraeus Medical Components) is an FDA-approved material. This demonstrated biocompatibility was one reason the PEDOT:PSS was selected for our study. Here, we measured the cytotoxicity of PEDOT:PSS NPs under relevant treatment conditions. We incubated murine epithelial cells (mIMCD-3) with PEDOT:PSS nanofluids of increasing concentrations (0.006 wt.% to 0.1 wt.%) for two durations (1 hour or 24 hours). The cell viabilities from all experiments are summarized in **Figure 5**. Only cells incubated in 0.1 wt.% PEDOT:PSS nanofluid exhibited a noticeable decrease in cell viability after 1 hour incubation. The viability of the cells started to decrease at a concentration higher than 0.05 wt.% for 24 hours of incubation. Considering the moderate concentration (::S 0.03 wt.%) and the typical duration of LL treatment (∼1 hour), the use of a PEDOT:PSS nanofluid in LL is promising.

**Figure 5.**
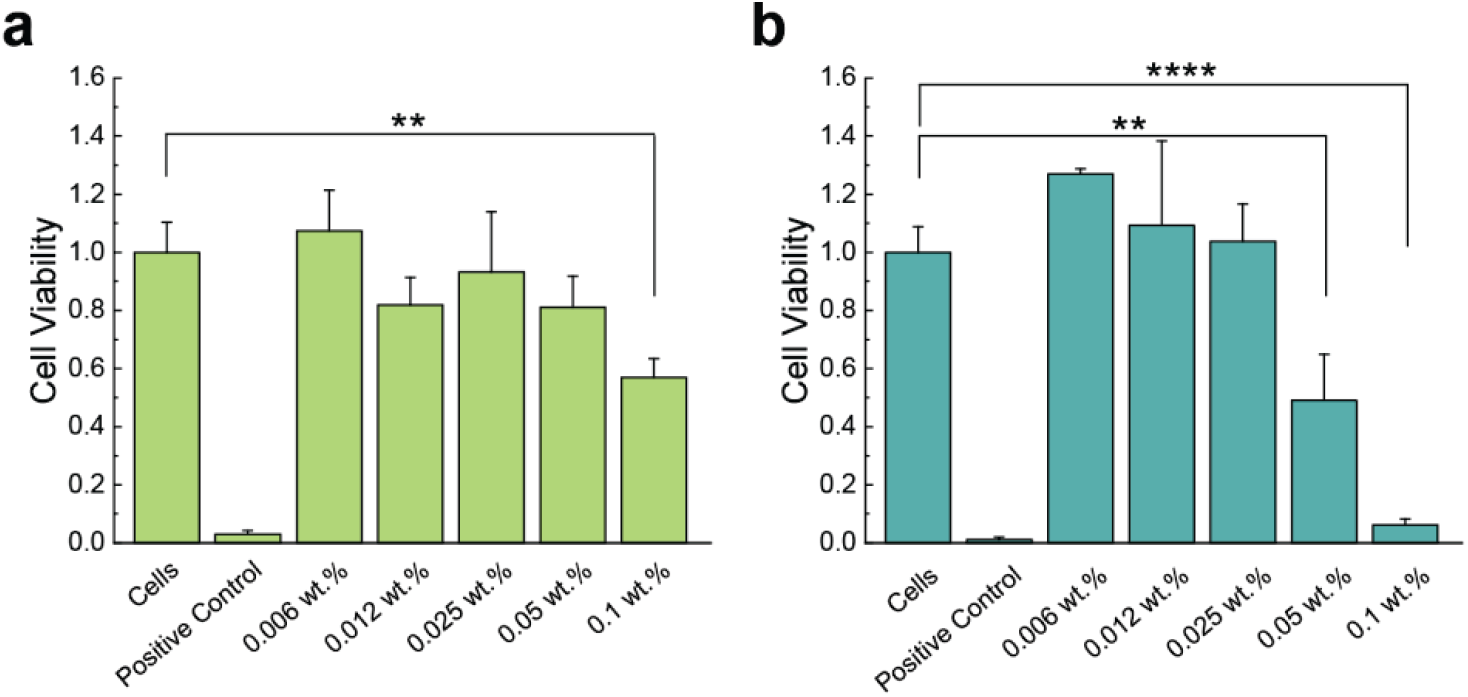
Viability of mIMCD-3 cells, measured with a MTT assay, after incubation with increasing concentrations of PEDOT:PSS. Exposure to triton X-100 (10% in PBS, 30 s) was used as a positive control to decrease cell viability. (a) 1 h incubation. (b) 24 h incubation. Significance was measured using 1-way ANOVA with Dunnett’s multiple comparisons post hoc. **p<0.01, ****p<0.0001, ns – not significant comparison are not presented.

## Conclusion

In this study, we presented the first demonstration of using a nanofluid to enhance the treatment of kidney stones. By incorporating PEDOT:PSS NPs into the fluid, we enhance the ablation efficiency of Ho:YAG LL. At a concentration of 0.03 wt.%, PEDOT:PSS solution exhibited a 25% increase in Ho:YAG laser absorption coefficient compared to water. For all three tested standoff distances, this solution demonstrated significant increase in stone ablation efficiency: over 38% improvement in spot treatment and over 26% improvement in the scanning treatment. The observed improvement in stone ablation was mainly attributed to two factors: (i) the rapid establishment of a vapor tunnel, and (ii) the enhanced laser energy absorption due to nanofluid infiltration. Aside from the increased absorbed laser energy accounting for the enhanced photothermal ablation, we believe that micro-explosions of the trapped PEDOT:PSS nanofluid inside the stone also contribute to the improved overall ablation efficiency.^[12,51,68]^ Micro-explosion is the phenomenon of fluid vaporization inside the stone pores that causes dramatic fracturing due to the high-pressure shockwaves. However, quantitative experimental and theoretical verification related to this contribution are complex and nontrivial and will be investigated in future studies. Last but not least, cell viability tests and temperature measurements show the nanofluid approach’s safety against cell damage.

Overall, PEDOT:PSS nanofluid has demonstrated significant potential for enhancing the efficiency of Ho:YAG LL in removing urinary stones. Our work distinguishes itself from previous studies by leveraging NIR absorber-induced efficiency improvements, marking a notable advancement in LL that can potentially shorten the operation time and patients’ burden without compromising laser safety.

## Materials and Methods

Aqueous dispersion of PEDOT:PSS conductive polymer (Clevios PH 1000) was purchased from MSE Supplies and used without further purification. The solution has a concentration of 1.2 wt.% and the average particle size is 30 nm. The solution was diluted by DI water into various concentrations. Artificial BegoStone phantoms (6×6 mm pre-soaked cylinders; 5:2 powder to water ratio, BEGO USA, Lincoln, RI, USA), a clinical Ho:YAG laser lithotripter (H Solvo 35-watt laser, Dornier MedTech, Munich, Germany), a flexible ureteroscope (Dornier AXISTM, with a 3.6 F working channel from Munich, Germany) with a 270 μm laser delivery fiber (Dornier SingleFlex 200, Munich, Germany) were used throughout this project.

Near-IR absorption spectroscopy: The NIR absorbance spectra of solutions were measured using a UV-Vis-NIR spectrophotometer (Cary 5000, Agilent, Santa Clara, CA, USA). During the measurement, the solution was filled in a cuvette with a path length of 0.2 mm (IR quartz, FireflySci, Inc. Northport, NY, USA) and the scan range was set to be 1350 – 2500 nm (resolution: 1 nm and scan rate: 600 nm/min). The slit width was 1 nm. An empty cuvette was used as the background when collecting data.

Visible transmission spectroscopy: The visible spectra were measured on the UV-Vis-NIR spectrophotometer (Cary 5000, Agilent, Santa Clara, CA, USA). The scan range was 380 – 750 nm (resolution: 1 nm and scan rate: 600 nm/min). The slit width was 1 nm. A glass cuvette with a path length of 5 mm filled with water was used as the background when collecting data. The average transmittance of a solution in visible is calculated by:

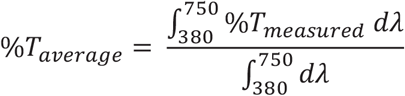

NIR diffusive reflectance: The NIR diffusive reflectance of the BegoStone samples was measured using a UV-Vis-NIR spectrometer (Shimadzu UV3600 Plus) equipped with an integrating sphere. The scan range was set to be 1950 – 2250 nm (resolution: 1 nm and scan rate: medium). The slit width was 1 nm. A standard white plate coated with BaSO_4_ was used as the reference. 50 μL of the fluid is dropped on the top surface of the BegoStone and it quickly diffused into the stone within 30 s and no liquid can be observed on the stone surface. For each fluid, the same measurement was repeated on seven different stones to account for the variation in stones. The average of seven measurements and standard deviation (colored shade) were plotted in Figure 3g. Since BegoStone is very thick (6 mm), we assume no light can transmit through, so stone absorption = 1 – reflectance.

LL in air: We used the same method to prepare the wet stones as in the NIR diffusive reflectance measurement. With the laser fiber (Dornier SingleFex 270, numerical aperture = 0.26) contacting the stone surface, laser pulses (60 pulses, n = 5) were delivered at an energy level of 0.2 J and a frequency of 20 Hz in dusting mode using a clinical Ho:YAG laser lithotripter (H Solvo 35-watt laser, Dornier MedTech). The resultant damage craters were then scanned by optical coherence tomography (OCT) to extract crater volumes, depths, and profile areas.^[34]^

Spot treatment in fluids: The spot treatment was conducted in a quartz cuvette by using the same Ho:YAG laser lithotripter. Within the cuvette, a thick layer of transparent hydrogel (Gelatin #1; Humimic Medical, SC, USA) was applied to its inner walls to simulate the soft boundary of kidney tissue, leaving a 12×12×40 mm^3^ cuboid space in the middle as depicted in Figure 3d. To further mimic the calyx environment of the kidney, a 3D-printed part with a spherical chamber of 5 mm in radius was fixed in the cuboid. The chamber was filled with 2 mL of PEDOT:PSS solutions at different concentrations (0 wt.%, 0.01 wt.%, and 0.03 wt.%) and the stone sample was positioned in its center. Laser pulses (60 pulses, n = 5) were delivered at an energy level of 0.2 J and a frequency of 20 Hz in dusting mode at various fiber tip-to-stone standoff distances (SDs; 0, 0.5, and 1.0 mm). The resultant damage craters were then scanned by optical coherence tomography (OCT) to extract crater volumes, depths and profile areas.

Scanning treatment in fluids: A BegoStone slab (23 x 23 x 10 mm^3^, L x W x H) was immersed in a large container filled with ∼ 20 mL of the fluid (water or 0.03 wt.% PEDOT:PSS nanofluid). The fiber integrated ureteroscope was mounted on a motorized stage with a fixed moving speed of 0.3 mm/s. Laser pulses (6000 pulses, n = 5) were delivered at an energy level of 0.2 J and a frequency of 20 Hz in dusting mode at various fiber tip-to-stone standoff distances (SDs; 0.1, 0.5, and 1.0 mm). After the treatment, the stones were dried in an oven (∼70°C) for 24 h and then sat on the benchtop for another 24 h. Afterwards, the dry stones were weighed on the balance. The difference between the initial and final masses were used to evaluate the treatment efficiency.

Temperature measurement in hydrogel kidney model: The hydrogel kidney model was prepared according to the method described in the literature.^[59]^ Seven K-type thermocouples (OMEGA, Norwalk, CT, USA) punched through the wall of the hydrogel and were in contact with the fluid in the upper calyx. Two halves of the model were clamped together and then submerged in the tank filled with room temperature water. The fiber-integrated ureterosope was inserted into the upper calyx through the opening on the side. Before turning on the laser, the upper calyx was flushed with the fluid (water or 0.03 wt.% PEDOT:PSS nanofluid) at room temperature for 15 s at a flow rate of 20 mL/min controlled by a peristaltic pump. The temperature was recorded with an interval of 0.2 s, and each experiment was repeated 5 times. To assess the potential risk of thermal injury, the thermal dose, evaluated by the cumulative equivalent minutes at 43 °C (CEM43°C), was calculated using:^[60]^

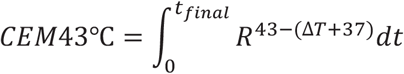

where R is 0.25 for T < 43°C and 0.5 for T ≥ 43°C.

Cell Culture: A murine epithelial cell line (mIMCD-3, ATCC, Manassas, VA, USA) was used for all experiments. Cells were cultured in Dulbecco’s Modified Eagle Medium/Hams F-12 (DMEM/F-12; pH 7.4, #11320033, Thermo Fisher, Waltham, MA, USA), supplemented with 10 % fetal bovine serum (FBS, #10437028, Thermo Fisher, Waltham, MA, USA). Cells grew at 5% CO_2_ at 37 °C and passaged upon reaching 80% confluency with a maximum passage number of 20.

MTT Assay: MIMCD-3 cells were seeded onto sterile 24-well plates (#3524, Corning Inc, Corning NY, USA) at a density of 150,000 cells per well. The cells were cultured overnight and incubated with PEDOT:PSS (0.006 wt.%-0.1 wt.%) for 1 h or 24 h. Control cells were incubated in DMEM/F-12 media for the duration of the treatment. Incubation with Triton X-100 (10% in Dulbecco’s Phosphate Buffered Saline (PBS), 30 s exposure; X100-100ML, Millipore Sigma, Burlington, MA, USA) was used to damage cells, serving as positive control. Cells were washed 3 times in PBS (#28374, Thermo Fisher, Waltham, MA, USA). Cells were incubated in DMEM (300 μl; #31053028, Thermo Fisher) and 3-(4,5-dimethylthiazol-2-yl)-2,5-diphenyl tetrazolium bromide (MTT, 0.5mg/ml, #V13154, Thermo Fisher, Waltham, MA, USA) for 1 h. Media was aspirated, and cells were incubated in dimethyl sulfoxide (300 μl; DMSO, #D8418, Millipore Sigma) for 10 minutes in the dark. DMSO was transferred into a clear 96-well plate (#82050, VWR, Radnor, PA, USA) and absorbance was measured at 580 nm using a plate reader (SpectraMax iD3, Molecular Devices, San Jose, CA, USA).

## Acknowledgement

The authors acknowledge the financial support from NIH National Institute of Diabetes and Digestive and Kidney Diseases (1P20-DK135107-01).

## Author contributions

P.-C. H. conceived the study and were responsible for project management. Q. F. performed the optical measurements on nanofluid. J. C. performed the stone damage assessment and collected bubble dynamics data. A. M. prepared the hydrogel kidney model and A. S. performed the LL in the kidney model. F. A. performed the cytotoxicity experiment. T.-H. C., J. D. and C. P. were involved in conceptualization of the study. Q. F. analyzed all the data and drafted the manuscript with P.-C. H. All other authors provided critical feedback and helped shape the study and edit the manuscript.

## Competing interests

P.-C. H., P. Z. and Q. F. have filed a patent application related to this work.

**Correspondence** and requests for materials should be addressed to Po-Chun Hsu and Pei Zhong.

## Notes

### Competing Interest Statement

The authors have declared no competing interest.

### Summary of Updates

New data of artificial kidney model experiments was added.

