## Supplemental information for "Nanofluid-Enhanced Laser Lithotripsy Using Conducting Polymer Nanoparticles"

### Note S1. Visibility test using the ureteroscope

The offset distance between the fiber tip and the ureteroscope was set to 5 mm in the visibility test and there was an additional 2 mm set for the SD, so the distance between the ureteroscope and the stone surface in the images below was 7 mm in total. The edge of the stone starts to be blurry at the concentration of 0.06 wt.% and it becomes totally indistinguishable when the concentration further increases into 0.12 wt.%.

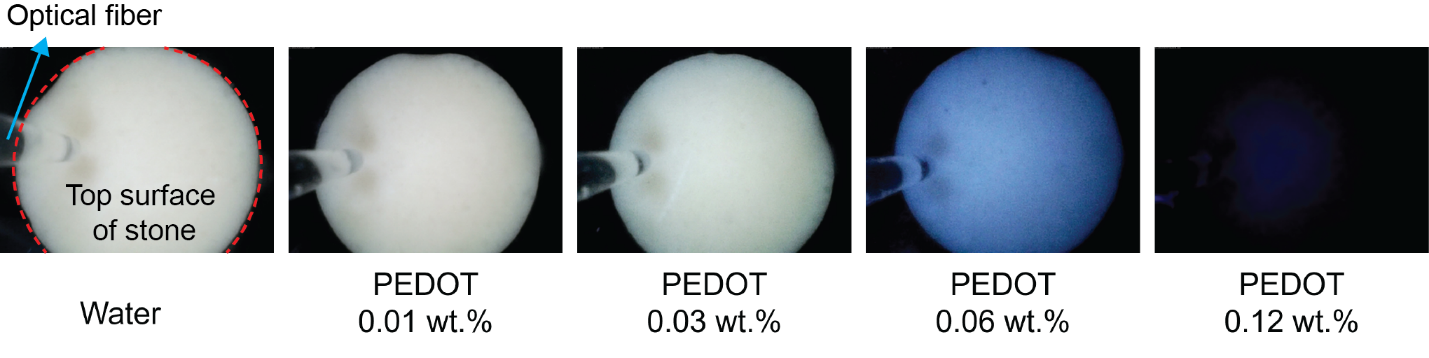

**Figure S1.** The appearance of the BegoStone under ureteroscope in PEDOT:PSS solutions with different concentrations. The red dashed line outlines the edge of the top surface of the BegoStone.

### Note S2. The measurement of the bubble width and distance between the bubble front and the fiber tip

The images captured by the high-speed video camera were imported in MATLAB for image processing to measure the distance between the bubble apex and the fiber tip. The original frames captured by the camera have a relative low resolution (112*104 pixel^2^) with coarse pixels, After the image was imported into MATLAB, a resize function was used to enlarge the image by 20 times (2240*2028 pixel^2^) with a “bilinear” interpolation method utilized to assign the intensity for the generated pixels. The actual distance of each pixel in the image was determined by a calibration photo of a ruler (Figure S2).

The measurement in MATLAB was done through recognizing the abrupt change in pixel brightness at the boundary. The position of the laser fiber tip was first located in the first frame where no vapor bubble is present to avoid the optical distortion it may cause.

The measured data of each curve in Figures 2c is the average and standard deviation of 45 measurements (the 20th, 25th, 30th, 35th, 40th, 45th, 50th, 55th and 60th pulses of 5 measurements) to minimize the variation caused by the laser output power shown in Figure S4.

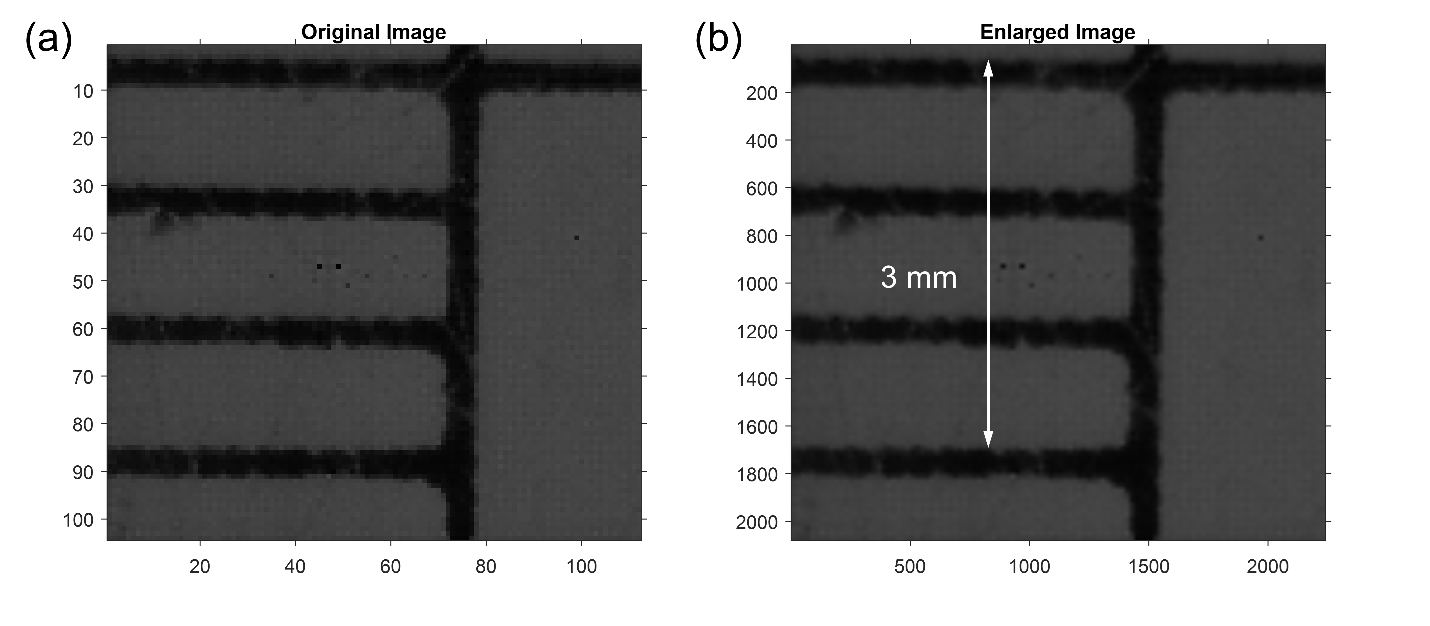

**Figure S2.** (a) Original image with low resolution. (b) Enlarged image with 20 times higher resolution. This image is also used to determine the distance of each pixel.

### Note S3. Measurement of surface tension of PEDOT:PSS solutions

Similar faster bubble expansion behavior was also observed in experiments where surfactants were added to water to reduce its surface tension.^[1,2]^ For example, Giglio et al.^[1]^ and Saeed et al.^[2]^ observed that a reduced surface tension leads to elongated bubble lengths and lifetimes in the fluid although the ablation efficiency of the stones under such conditions was not explored. However, unlike surfactants with amphiphilic molecular structures, PEDOT:PSS NPs have a hydrophilic shell that enables their dispersity in water.^[3]^ Therefore, when these NPs are dispersed in water, they remain inside the water instead of at the water-air interface, resulting in no change to the surface tension. To confirm this, we measured the surface tension of PEDOT:PSS nanofluids with various concentrations using the automatic surface tensiometer (BZY-201, Shanghai Fangrui Instrument Co., Ltd, Shanghai, China). Five independent measurements were made for each solution, and the average values were plotted in Figure S3.

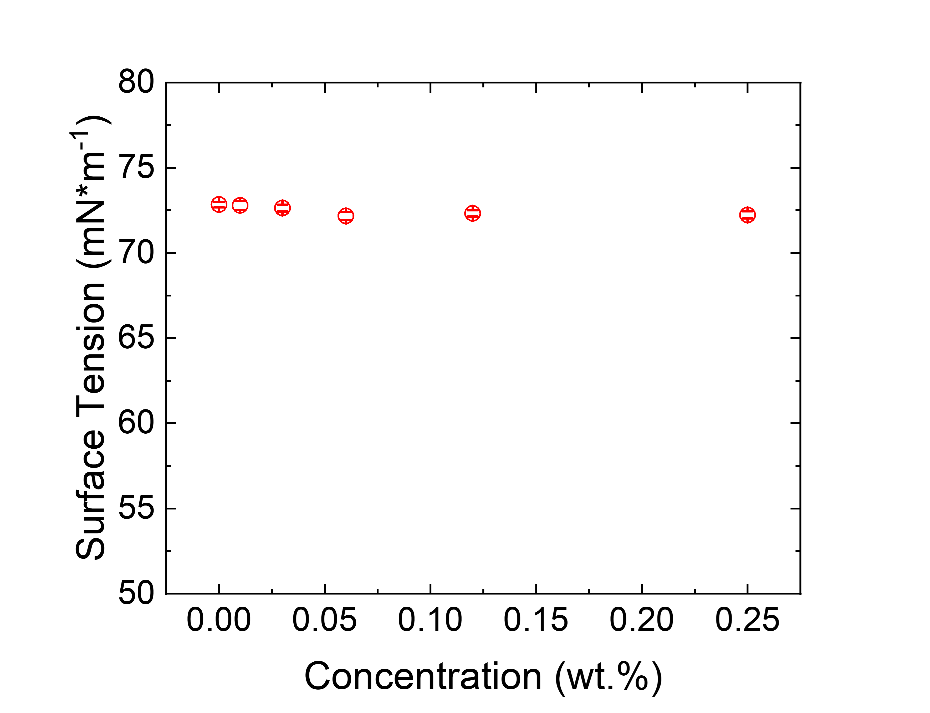

**Figure S3.** Surface tension of H_2_O and PEDOT:PSS solutions with different concentrations.

### Note S4. The calculation of temporal transmittance of 2120 nm light

Time-dependent transmittance is calculated based on the Beer-Lambert law by considering the distance shown in Figure 2c:

$$T\left( t \right)=\frac{I(t)}{I_{0}}=e^{-\alpha_{v}l_{v}(t)-\alpha_{l}l_{l}(t)}$$

where $\alpha_{v}$ and $\alpha_{l}$ are the absorption coefficient of the vapor and liquid phase. $l_{v}$ and $l_{l}$ are the light path in the vapor and liquid phase from the fiber tip to the detector at a specific standoff distance. The sum of $l_{v}$ and $l_{l}$ was equal to the SD (0.5 mm or 1 mm). Table S1 listed all the values we used in this calculation.

|  | **Absorbance@2120 nm** | **Abs. Coeff. @2120 nm (mm^-1^)** |
| --- | --- | --- |
| Vapor |  | 0.001 |
| Water | 0.22885 | 2.63522 |
| 0.01 wt.% PEDOT | 0.25389 | 2.92356 |
| 0.03 wt.% PDEOT | 0.30462 | 3.50775 |

**Table S1.** Measured Absorbance ($Absorbance= -\log_{10} T$) and derived absorption coefficients of different fluids to the Ho:YAG laser. (Abs. Coeff. = 2.303*Absorbance/path length, where path length = 0.2 mm).

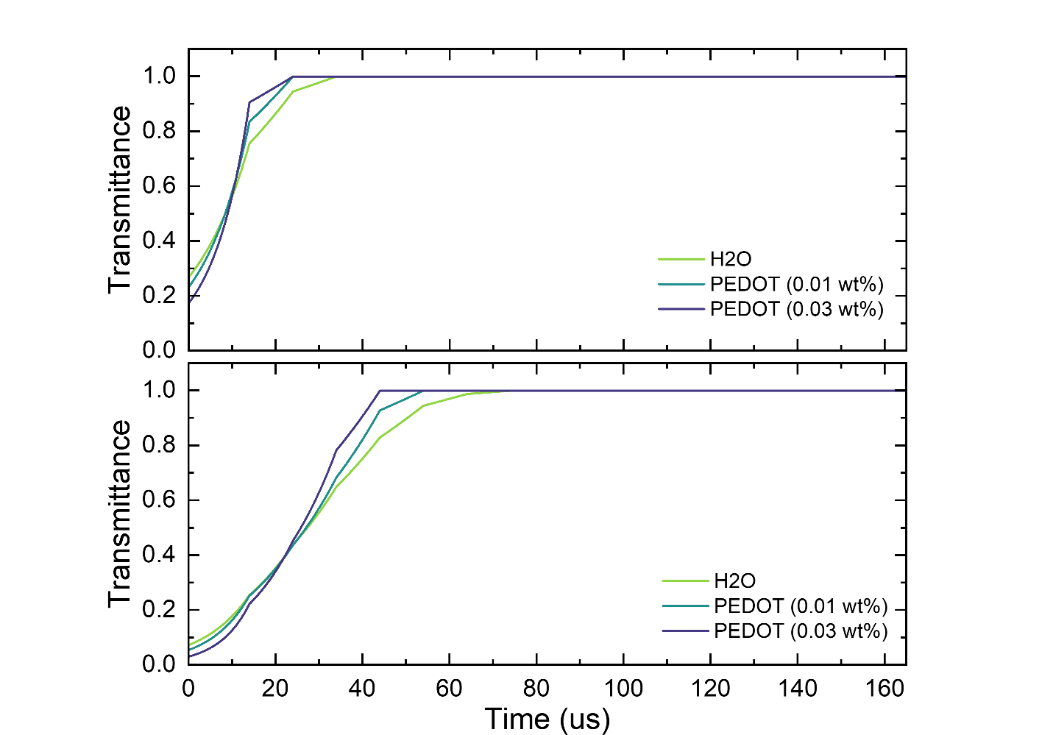

**Figure S4.** Calculated time-dependent transmittance of 2120 nm light in different fluids at the SD = 0.5 mm (top) and 1 mm (bottom).

### Note S4. Laser output power measurement

The laser output profile was measured using a photodiode.^[4]^ The laser delivery fiber was submerged in water and positioned perpendicularly to a 1 mm thick glass slide. The fiber tip was contacting the glass slide, and a light guide was positioned approximately 20 mm away on the opposite side of the glass slide in air to collect the transmitted light to an InGaAs photodetector (PDA10D, Thorlabs, Newton, NJ). All 60 pulses were recorded in each measurement and the experiments were repeated independently five times. Figure S5a shows that the laser output not only fluctuates within each measurement, but also shows variance among different repeats. To minimize the error caused by the random fluctuation of laser output, the average laser output of a total 45 pulses (the 20^th^, 25^th^, 30^th^, 35^th^, 40^th^, 45^th^, 50^th^, 55^th^ and 60^th^ pulses of each measurement) was taken to be 0.2 J. Figure S5b is the laser output profile of the 60^th^ pulses in 5 measurements.

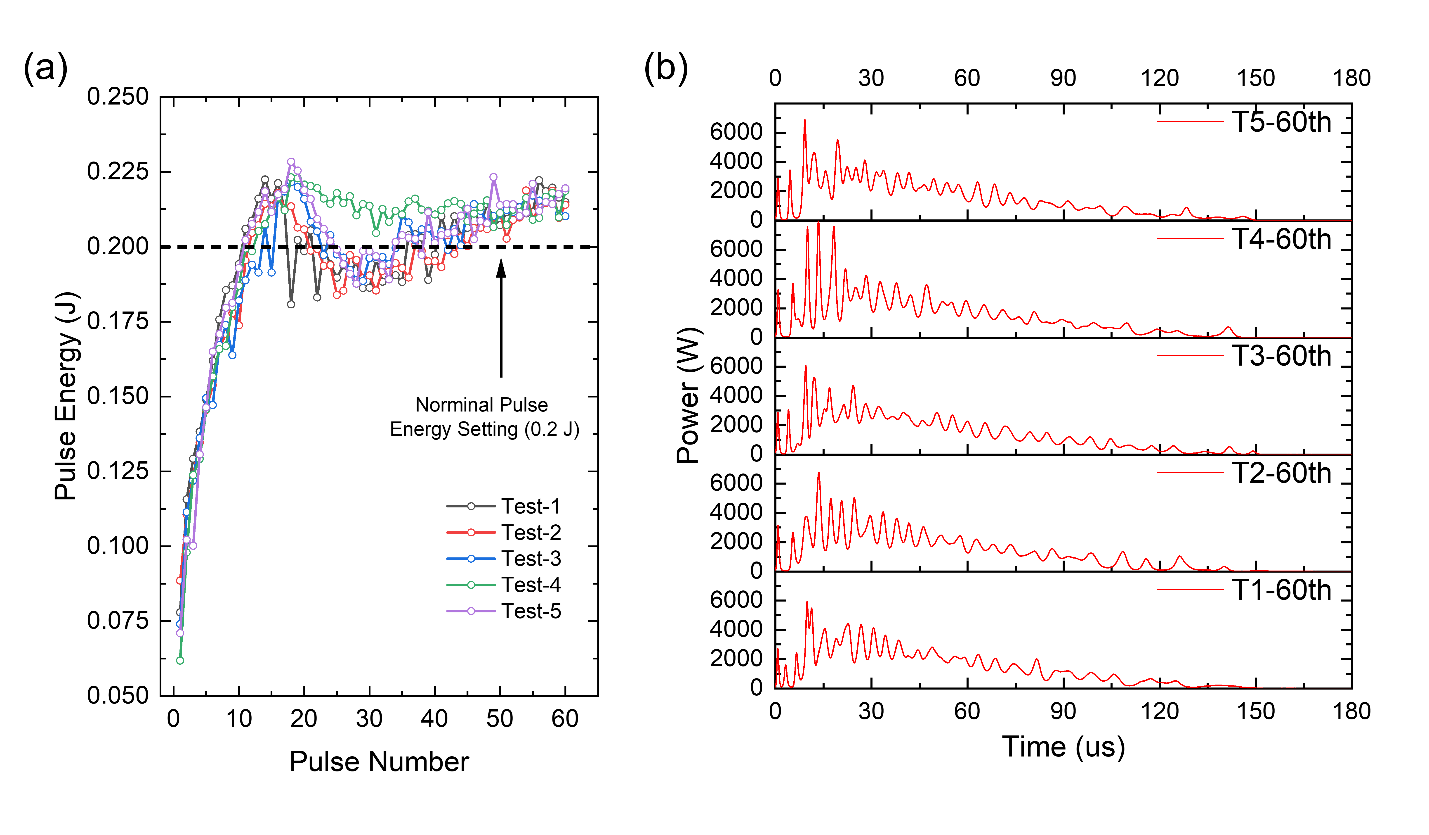

**Figure S5.** (a) The pulse energy delivered by the Ho:YAG laser. (b) The laser output profile of the 60^th^ pulses in 5 measurements.

### Note S5. Calculation of transmitted laser energy

The energy transmitted to the stone surface at a specific standoff distance can be calculated using:

$$Delivered energy=\int_{t=0}^{t=160 \mu s} P_{laser source}*Tdt$$

Laser output of 45 pulses (the 20^th^, 25^th^, 30^th^, 35^th^, 40^th^, 45^th^, 50^th^, 55^th^ and 60^th^ pulses of 5 independent measurements) were applied to this equation separately and its average and standard deviation were summarized in Figure 3e.

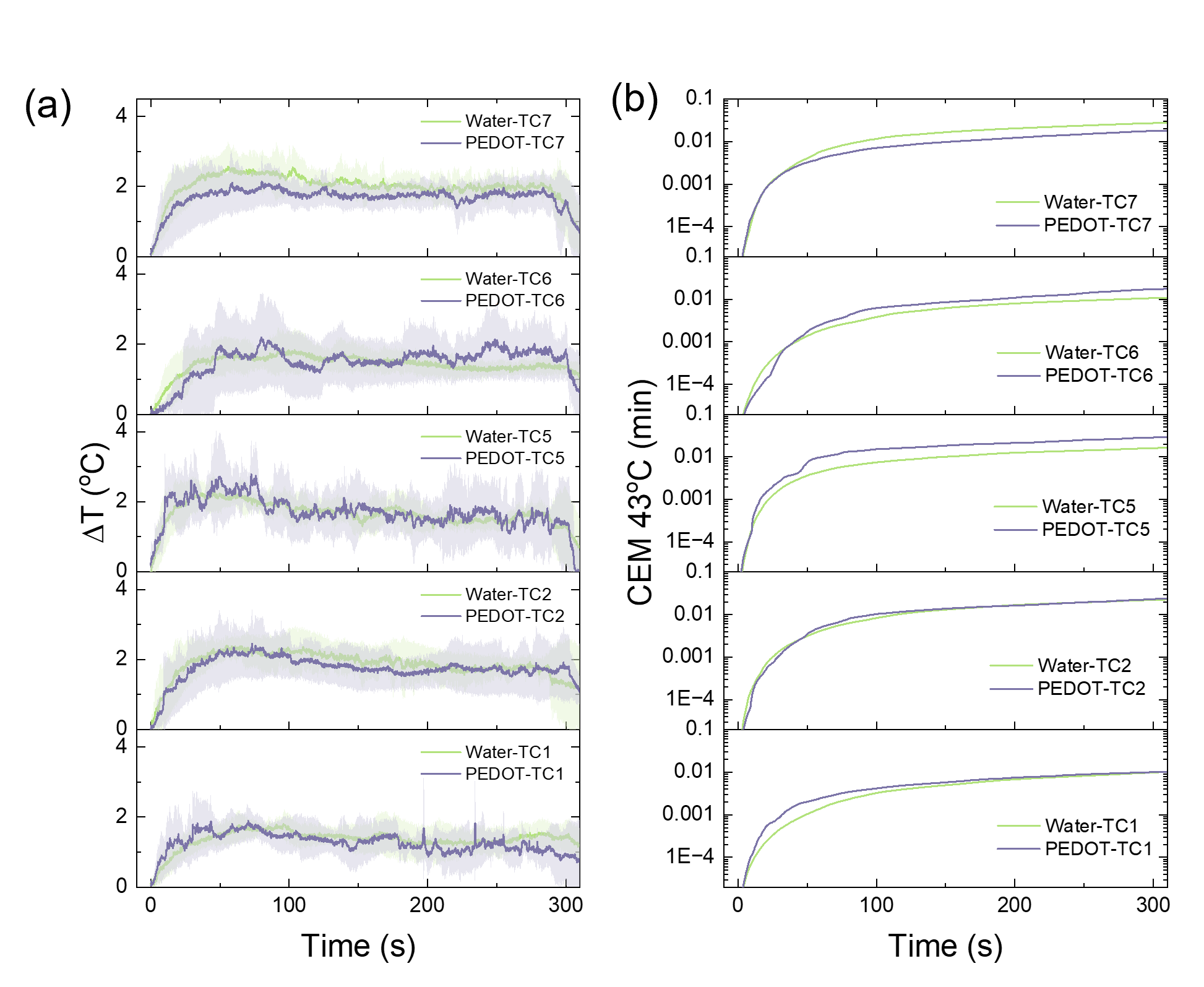

**Figure S6.** (a) The temperature changes of thermocouples #1, #2, #3, #5, #6 and #7 during the LL. The solid line and shaded areas are the mean and standard deviation of 5 independent measurements. (b) The corresponding CEM 43°C calculated based on the average temperature change in (a).

### Note S6. PEDOT:PSS nanoparticles in saline

In clinical practice, the urologists irrigate the patient’s kidney/bladder/ureter using saline (0.9 wt.% NaCl aqueous solution) to prevent the cell damage due to the imbalanced osmotic pressure. In recognizing this, we also explored the performance of PEDOT:PSS nanoparticles dispersed in saline via the spot treatment (Ep = 0.2 J, F = 20 Hz, 60 pulses, SD = 0 mm). We noticed that there were some large aggregates of PEDOT:PSS particles in saline. This is because of the partial screening of the electrostatic repulsion among the nanoparticles at such a high ionic strength condition. Aggregates with size larger than microns cannot diffuse into the pores in the BegoStone, therefore diminishing the efficiency of PEDOT:PSS nanofluid from 114% to 71%. However, such an issue may be resolved by increasing the charge of the nanoparticles via surface coating/modification, which will be investigated in the future.

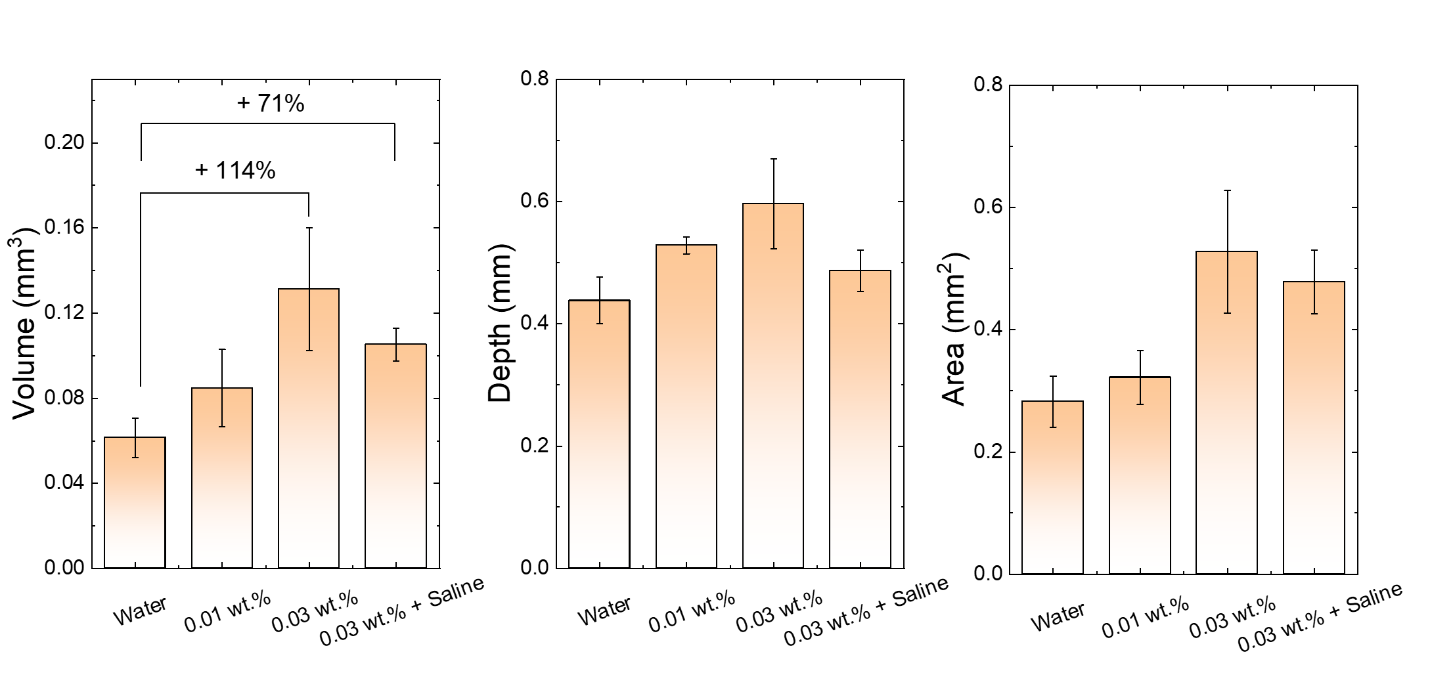

**Figure S7.** Volume, maximum depth and profile area of the craters produced on BegoStones via spot treatment. 0.01 and 0.03 wt.% PEDOT:PSS nanofluid was diluted using water and 0.03 wt.% + saline was diluted using 0.9 wt.% NaCl solution.
